# Development of subunit selective proteasome substrates for *Schistosoma species*

**DOI:** 10.1101/2024.02.13.580161

**Authors:** Zhenze Jiang, Elany B. Silva, Chenxi Liu, Pavla Fajtová, Lawrence J. Liu, Nelly El-Sakkary, Danielle E. Skinner, Ali Syed, Steven C Wang, Conor R. Caffrey, Anthony J. O’Donoghue

## Abstract

Schistosomiasis, or bilharzia, is a neglected tropical disease caused by *Schistosoma* spp. blood flukes that infects over 200 million people worldwide. Just one partially effective drug is available, and new drugs and drug targets would be welcome. The 20S proteasome is a validated drug target for many parasitic infections, including those caused by *Plasmodium* and *Leishmania*. We previously showed that anticancer proteasome inhibitors that act through the *Schistosoma mansoni* 20S proteasome (Sm20S) kill the parasite *in vitro*. To advance these initial findings, we employed Multiplex Substrate Profiling by Mass Spectrometry (MSP-MS) to define the substrate cleavage specificities of the three catalytic β subunits of purified Sm20S. The profiles in turn were used to design and synthesize subunit-specific optimized substrates that performed two to eight fold better than the equivalent substrates used to measure the activity of the constitutive human proteasome (c20S). These specific substrates also eliminated the need to purify Sm20S from parasite extracts - a single step enrichment was sufficient to accurately measure substrate hydrolysis and its inhibition with proteasome inhibitors. Finally, we show that the substrate and inhibition profiles for the 20S proteasome from the three medically important schistosome species are similar, suggesting that data arising from an inhibitor development campaign that focuses on Sm20S can be extrapolated to the other two targets with consequent time and cost savings.

## Introduction

Schistosomiasis, or bilharzia, is a neglected tropical disease caused by *Schistosoma* spp. blood flukes. The disease thrives in areas that lack access to safe drinking water and adequate sanitation, and the World Health Organization estimates that 251.4 million are in need of treatment. *Schistosoma mansoni*, *S. haematobium* and *S. japonicum* are the principal etiological agents of disease in humans. Adult worms in the bloodstream lay eggs that lodge in various visceral organs, including the spleen, liver, intestine and bladder. The eggs induce chronic inflammatory and fibrotic reactions that are responsible for malaise, pain and disability, which means lost school days for children and reduced worker productivity ^1–3^. Thus, schistosomiasis, as a chronic and morbid disease, is as much a contributor to poverty as it is a consequence of it.

Since the early 1980s, treatment and control of schistosomiasis has relied on one drug, praziquantel (PZQ) ^4–6^, which is primarily delivered via mass drug administration programs that aim to reduce disease morbidity at the community level^7, 8^. Although reasonably effective at the single oral dose offered ^9–12^, is much less active against developing worms ^9, 13–16^, which after treatment, mature and continue to generate morbidity. In addition, and apart from the ever-present threat of drug resistance^9, 17, 18^, PZQ has a number of pharmaceutical and pharmacological problems^5, 6, 9, 19^ that encourage the search for new drugs.

Among the potential molecular drug targets discussed over the years for chemotherapy of schistosomiasis^6, 20^, a recent addition has been the proteasome^21, 22^ which is an evolutionarily conserved, multi-subunit, ATP-dependent proteolytic complex that is vital to cellular proteostasis ^23, 24^. Each 20S proteasome core comprises two stacked rings of seven β subunits between two rings of seven α subunits (**Fig. 1A**). Among the seven β subunits in each ring, three subunits (β1, β2 and β5) are proteolytically active, and, based on their peptidyl cleavage specificities, are described as having caspase-, trypsin- and chymotrypsin-like specificities, respectively. The human constitutive 20S proteasome (c20S) is a well-established target for various cancers, and the Food and Drug Administration has approved three small molecule inhibitors, bortezomib (BTZ), carfilzomib (CFZ) and ixazomib (IXZ). These drugs preferentially target β5; however, at higher concentrations, BTZ and IXZ also inhibit β1, and CFZ inhibits β2. The marine natural product marizomib (MZB) is a brain-penetrable proteasome inhibitor that preferentially targets the β5 subunit, but also inhibits β1 and β2 at higher concentrations ^25^. MZB is in phase 3 clinical trials for the treatment of glioblastoma ^26^.

**Figure 1.**
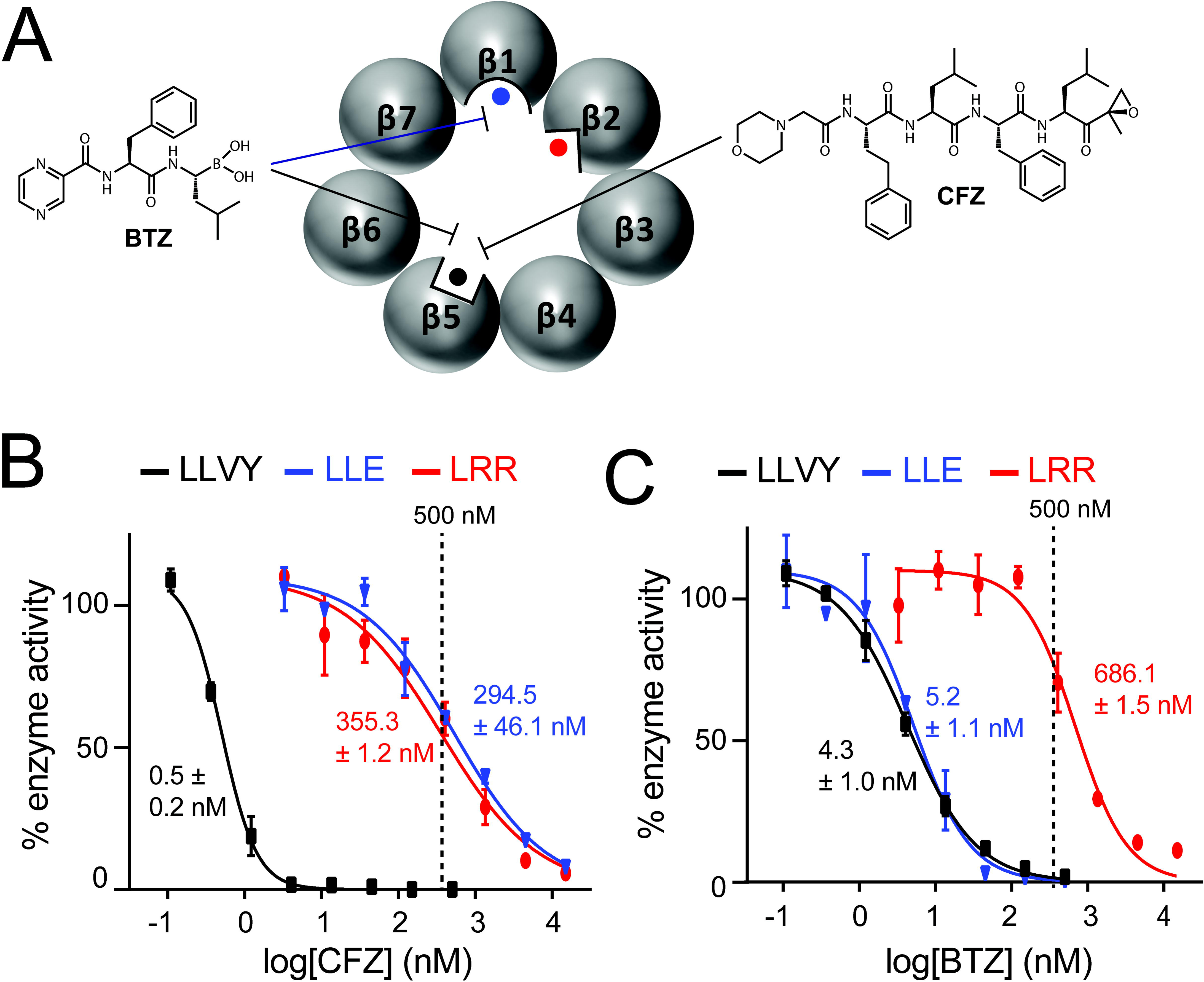
Determining the inhibitor concentration needed to assay individual Sm20S β subunit activity. **A**. Schematic representation of the β5 ring of Sm20S showing the β1, β2, and β5 catalytic subunits with differently shaped active sites to illustrate differences in inhibitor specificity. Also shown are the chemical structures of CFZ and BTZ, and the Sm20S subunits that each preferentially inhibits. **B, C**. The potency of CFZ and BTZ was determined for each catalytic subunit using Z-LLE-amc for β1, Z-LRR-amc for β2, and Suc-LLVY-amc for β5. Assays were performed in technical triplicates.

In spite of its evolutionary conservation, active site differences between the human and parasite proteasomes have been identified and exploited to develop inhibitors that specifically target the latter ^21, 27–36^. For example, researchers have discovered potent inhibitors of the *Plasmodium falciparum* proteasome (Pf20S) based on either cleavage profiling of peptidyl substrates^27^ or the screening of proteasome inhibitor libraries ^37–39^. A selection of covalent inhibitors containing epoxyketone ^28, 40^, vinyl sulfone ^41^ and boronic acid reactive groups (warheads) ^39^ with selectivity indices ranging from 56 to 2,640 between blood stage parasites and human cell lines have been reported. Our own research program to develop Pf20S inhibitors has synthesized over 150 analogs of the marine natural product, carmaphycin B ^21, 28, 42–44^, which contains the epoxyketone reactive group. One of these demonstrated a proof-of-concept therapeutic benefit in a mouse model of *Plasmodium* infection ^35, 40^. In addition to malaria, various groups have shown that proteasome inhibitors are effective agents in animal infection models of the kinetoplastid parasites, *Trypanosoma brucei*, *Trypanosoma cruzi* and *Leishmania donovani.* ^32, 35, 45^ In one striking case of convergent drug discovery, four groups independently identified the azabenzoxazole chemotype that offered low nanomolar killing of the respective kinetoplastid parasites *in vitro* and cured infection in the respective animal models ^32, 45–48^. Two drug candidates to emerge from this line of research, GSK245 (GSK) and LXE408 (Novartis) are now in Phase I and II clinical trials respectively, for treatment of visceral leishmaniasis caused by *L. donovani* ^33, 49^. Of note, both compounds bind to the interface of the β4 and β5 subunits that is unique to the trypanosomatid 20S to sterically block proteolytic activity ^32^.

We previously demonstrated that targeting the *Schistosoma mansoni* 20S (Sm20S) *in vitro* with proteasome inhibitor drugs approved for cancer treatment leads to immobility and, ultimately, fatal degenerative changes in the adult worm ^21^. In addition, using a small collection of carmaphycin B analogs, we identified one Sm20S inhibitor that possessed a 27-fold improved selectivity index *vs*. HepG2 cells, suggesting that a path forward exists to build carmaphycin B-based inhibitors with greater specificity for the Sm20S compared to the c20S anti-target.

Here, we describe the purification of Sm20S and define its substrate cleavage specificities using Multiplex Substrate Profiling by Mass Spectrometry (MSP-MS) ^50^. The profiling data arising were used to design fluorogenic peptidyl substrates specific to each of the three proteolytically active β subunits. We show that each new substrate is more efficiently cleaved by Sm20S compared to those c20S substrates we previously used. Part of this process involved developing a method to selectively measure Sm20S proteolysis in extracts of *S. mansoni*. Last, we demonstrate that the cleavage profiles with the new substrates is similar across the three schistosome species most responsible for disease in humans. This suggests that a single campaign of inhibitor development focusing on Sm20S may be sufficient to develop a pan-schistosomicide, which would be consistent with the target product profile for schistosomiasis ^51, 52^.

## Methods

### Preparation of schistosomes

The acquisition, preparation, and *in vitro* maintenance of *S. mansoni* have been described ^53, 54^. The Naval Medical Research Institute (NMRI) isolate of *S. mansoni* was cycled between *Biomphalaria glabrata* snails and male Golden Syrian hamsters (infected at 4-6 weeks of age) as intermediate and definitive hosts, respectively. In brief, adult schistosomes were harvested from hamsters 42 days post-infection in RPMI or DMEM, and washed five times prior to maintenance overnight at 37°C and 5% CO_2_ in Basch medium ^55^ containing 4% heat-inactivated FBS, 500 μg/mL streptomycin and 500 U/mL penicillin. The use of small vertebrate animals was approved by the Institutional Animal Care and Use Committee (IACUC) of the University of California San Diego. UCSD-IACUC derives its authority for these activities from the United States Public Health Service (PHS) Policy on Humane Care and Use of Laboratory Animals, and the Animal Welfare Act and Regulations (AWAR). Frozen *S. haematobium* and S*. japonicum* adult worms were shipped *gratis* by the Schistosomiasis Resource Center and the Biomedical Research Institute.

### Purification of the *S. mansoni* proteasome

Adult mixed-sex *S. mansoni* worms were washed and homogenized using a motorized Teflon pestle connected and 1.5 mL tubes containing ice-cold 100 mM Tris-HCl, 100 µM E-64, pH 7.5. Lysates were centrifuged for 15 min at 14,000×*g* and 4°C, and the supernatant subjected to two ammonium sulfate precipitation steps on ice at 30 and 60% saturation, respectively, each for 60 min. After centrifugation for 15 min at 14,000×*g* and 4°C, the supernatant was discarded and the precipitated proteins were resuspended in ice cold 100 mM Tris-HCl, 100 µM E-64, pH 7.5. The samples were enriched for Sm20S when concentrated using 100 kDa centrifugal filter units (Amicon). The protein concentration was quantified using the Pierce BCA kit.

Samples were either used for enzyme assays (defined as Sm20S-enriched) or subjected to column chromatography for Sm20S purification. For the latter, samples were loaded onto a Superose 6 10/300 gel filtration column under the control of an ÄKTA Pure instrument (GE Healthcare Life Sciences). Proteins were eluted using 50 mM HEPES, 10% glycerol, 0.125 M NaCl, pH 7.5. Fractions of 0.5 mL were collected and assayed for proteasome activity with 25 μM Succinyl-Leu-Leu-Val-Tyr-7-amino-4-methylcoumarin (Suc-LLVY-amc) in assay buffer (50 mM Tris, 0.02% SDS, pH 7.5). The assay was performed at 24°C using a Synergy HTX multi-mode reader (ex/em = 360 nm/460 nm; BioTek Instruments). Fractions containing proteasome activity were pooled and loaded onto a 5 mL anion exchange HiTrap DEAE FF column (Sigma). Proteins were eluted using 50 mM HEPES, 10% glycerol, pH 7.5, and a linear gradient of 0.125 to 0.6 M NaCl. Fractions (1.5 mL) were collected and assayed with Suc-LLVY-amc as described above. Fractions containing proteasome activity were pooled, concentrated using a 100 kDa centrifugal filter unit (Amicon) and stored at -80°C.

### Dose-response assays

Sm20S was preincubated for 15 minutes with either 20 μM to 9 nM or 740 nM to 0.3 nM of CFZ or BTZ and then assayed with 10 μM of Suc-LLVY-amc, Z-LLE-amc or Z-LRR-amc. The maximun velocity over 8 sequential readings was recorded and activity (RFU/sec) was normalized to the activity in the absence of CFZ or BTZ (0.1% DMSO). Dose response curves were generated using GraphPad Prism (version 10.0.0).

### Proteasome MSP-MS

MSP-MS was performed with a library of 228 synthetic tetradecapeptides that were designed to contain all neighbor and near-neighbor cleavage sites ^50, 56^. Sm20S was first incubated with 500 nM BTZ, CFZ or 0.1% DMSO (vehicle control). Assays were conducted in quadruplicate by incubating Sm20S with the peptide library at a final concentration of 0.5 µM for each peptide in 50 mM Tris-HCl, pH 7.5. Assays were incubated at 25°C for 15, 60 and 240 min. At each time point, 20 µL of the reaction mixture were removed and quenched by the addition of 80 μL 8M GuHCl. Samples were immediately stored at -80°C. Control reactions consisted of Sm20S pre-incubated with GuHCl to inactivate the enzyme prior to the addition of the peptide library. All samples were desalted using custom-made C18 spin tips and dried in a vacuum centrifuge. Samples were resuspended in 40 μL 0.1% formic acid, and ∼0.4 µg peptides were injected into a Q-Exactive Mass Spectrometer equipped with an Ultimate 3000 HPLC (Thermo). Peptides were separated by reverse phase chromatography on a C18 column (1.7 µm bead size; 75 µm x 25 cm; 65°C) at a flow rate of 300 nL/min using a 60-min linear gradient of 5 to 30% solvent B (0.1% formic acid in acetonitrile), with solvent A being 0.1% formic acid in water. Survey scans were recorded over a 150–2000 m/z range (70,000 resolutions at 200 m/z, AGC target 3×106, 100 ms maximum). MS/MS was performed in a data-dependent acquisition mode with HCD fragmentation (28 normalized collision energy) on the 12 most intense precursor ions (17,500 resolutions at 200 m/z, AGC target 1×10^5^, 50 ms maximum, dynamic exclusion 20 s). Data were processed using PEAKS 8.5 (Bioinformatics Solutions Inc.). MS2 data were searched against the tetradecapeptide library sequences with decoy sequences in reverse order. A precursor tolerance of 20 ppm and 0.01 Da for MS2 fragments was defined. No protease digestion was specified. Data were filtered to a 1% peptide level false discovery rate with the target-decoy strategy. Peptides were quantified with label free quantification, and data were normalized by median and filtered by 0.3 peptide quality. Missing and zero values were imputed with random normally distributed numbers in the range of the average of the smallest 5% of the data ± SD.

Cleaved peptides were identified in each dataset and the P4 to P4′ amino acids were inputted into iceLogo software and compared to all possible P4 to P4′ sequence in the library. Icelogo plots were generated that illustrate the frequency of amino acids present around the cleavage site.

### Proteasome activity and inhibition assays

Subunit specific fluorogenic substrates were custom synthesized and purified by HPLC to >95% by GenScript (New Jersey). Substrates contained either an N-terminal acetylation group and a C-terminal amc group, or an N-terminal 7-methoxycoumarin (mca) and a C-terminal lysine 2,4-dinitrophenyl (K(dnp)). Fluorogenic activity assays were performed in black, round-bottomed 96-well plates. Protein extracts from *Schistosoma* sp. were incubated with inhibitors for 1 h at room temperature and activity was assayed in a 50 µL total volume containing 7.5 µg of protein and 25 µM Suc-LLVY-amc in 20 mM Tris-HCl, 10 µM E-64, 0.02% SDS, pH 7.5. For purified Sm20S assays, 10 nM of enzyme was incubated with a 2-fold serial dilution of substrate. Controls contained 0.0001% DMSO. Fluorescence was monitored at 24°C in a Synergy HTX multi-mode reader (BioTek Instruments, Winooski, VT). Excitation and emission wavelengths for the amc and mca substrates were 360 and 460 nm, and 320 nm and 400 nm, respectively. Protease activity was quantified as RFU min^-1^ µg^-1^ protein and normalized to DMSO control reactions.

### Detection of 20S proteasome activity in three *Schistosoma* species using a fluorescent probe

*S. mansoni*, *S. haematobium* and *S. japonicum* protein extracts were subjected to ammonium sulfate precipitation as outlined above. The proteasome-containing fraction was concentrated and dialyzed in buffer containing 100 mM Tris-HCl pH 7.5, 50 μM E-64, 2 mM AEBSF, and 2 μM pepstatin using a 100 kDa centrifugal filter units (Amicon). Protein was quantified using the bicinchoninic acid assay and 2, 5, and 10 μg of protein were incubated with 2 μM of the activity-based probe, Me4BodipyFL-Ahx3Leu3VS (R&D Systems, #I-190) ^57^ at 37°C for 3 h. Samples were mixed with 4X NuPage lithium dodecyl sulfate loading buffer (Invitrogen), heated to 90°C for 5 min and then loaded onto 12% NuPAGE Bis-Tris Gels (Invitrogen). Proteins were separated using NuPAGE MOPS (3-(*N*-morpholino)propanesulfonic acid) as the running buffer. Human constitutive c20S at 20 nM was used as a positive control. Direct in-gel visualization of Me4BodipyFL-Ahx3Leu3VS-labeled proteasome subunits was explored using a fluorescence scanner with an emission filter at 530 nm. Based on the intensity of protein bands on the gel, a follow-up gel was run that contained 2.5 μg of *S. haematobium,* and 5 μg of *S. japonicum* and *S. mansoni* protein.

### Proteasome activity and inhibition assays in three *Schistosoma* species

Extracts of *S. haematobium* (0.625 μg/μL), or *S. japonicum* and *S. mansoni* (each 1.25 μg/μL) were enriched for proteasome by ammonium sulfate precipitation and the pre-incubated for 1 h with 500 nM BTZ, CFZ or MZB. In addition, the carmaphycin B analog, 17 ^21^ was preincubated at 100 nM, 500 nM or 2 μM with these samples. DMSO (0.125%) was used as a vehicle control. Proteolytic activity was quantified following the addition of an equal volume of 20 μM mca-VDQMDGW-K(dnp)-NH_2_ (β1-substrate) or mca-FnKRR-K(dnp)-NH_2_ (β2-substrate), or 40 μM Z-FNKL-amc (β5-substrate). Assays were performed in buffer containing 100 mM Tris-HCl pH 7.5, 50 μM E-64, 1 mM AEBSF, 2 μM pepstatin, and 0.03% SDS in a total volume of 8 μL in a 384-well plate (Greiner Bio-One #784900). The release of the fluorophore was recorded in a Synergy HTX plate reader with excitation at 340 nm and emission at 460 nm for the β5-substrate and excitation at 320 nm, emission at 400 nm for β1 and β2 substrates. Experiments were performed in triplicate wells, and DMSO was used as the negative control. The assay was performed at 24°C using a Synergy HTX multi-mode reader with excitation and emission wavelengths of 360 and 460 nm, and 320 nm and 400 nm, for the amc and mca substrates, respectively.

## Results

### Revealing the substrate specificities of *Schistosoma mansoni* proteasome subunits

We previously isolated the 20S proteasome from *S. mansoni* (Sm20S) and discovered inhibitors that target the β2 and β5 subunits. ^21^ To generate those data, we repurposed three human proteasome substrates, Z-LLE-amc, Z-LRR-amc and Suc-LLVY-amc that detect the activity of human β1, β2 and β5 subunits, respectively. However, when comparing the turnover rate of these substrates by c20S and Sm20S, it was evident that they were more efficiently cleaved by c20S. ^21^ This encouraged us to develop Sm20S subunit-specific substrates that would be cleaved with higher efficiencies and which would serve as valuable tools to screen for new inhibitors. Therefore, we utilized MSP-MS to define the cleavage specificities of each of the three Sm20S β subunits to then aid the development of new subunit specific substrates.

We have previously shown that c20S, the human 20S immunoproteasome (i20S) and Pf20S each cleave at more than 280 sites within the MSP-MS peptide library.^27, 58^ In this study, we isolated and incubated Sm20S with the same peptide library for 3 h and detected cleavage at 302 sites. We generated a specificity profile of this activity to show that Sm20S has a general preference for cleavage of peptides on the C-terminal side of Arg and Phe. However, it is impossible to know which subunit is responsible for cleaving each of these peptides. Physical separation of the 20S catalytic subunits is not possible without inactivation of the whole complex and so we used one or more inhibitors to selectively inactivate one or more subunits of Sm20S, such that the remaining non-inhibited subunit(s) could be assayed. To do this, we generated a dose-response curve with CFZ and determined that the compound inhibited the β5 subunit more potently than the β1 and β2 subunits. In the presence of 500 nM CFZ, the β5 subunit activity measured using Suc-LLVY-amc was completely inhibited, whereas the activities of β1 (Z-LLE-amc) and β2 (Z-LRR-amc) were only inhibited ∼40% (**Fig. 1B**). Therefore, we used 500 nM CFZ for follow-up MSP-MS studies.

Next, we performed a dose response study with BTZ and showed that the β5 and β1 subunits were preferentially inhibited over β2. Using 500 nM of BTZ, β5 and β1 were completely inactivated while activity of the β2 subunit was reduced by ∼35% (**Fig. 1C**). Thus, the data reveal that incubation of Sm20S with 500 nM CFZ will result in the β1 and β2 retaining activity while β5 is not active; while 500 nM BTZ will result in only β2 being active. Using these inhibitor-treated conditions, we then performed the MSP-MS assay and determined which subunits are responsible for cleaving which peptides. In the presence of 500 nM CFZ we anticipate that we could distinguish β5 from β1/β2 while 500 nM BTZ will distinguish β2 from β5/β1. By a process of elimination, we will be able to determine the specificity of β1 as being the activity that is not β2 or β5.

### Development of a Sm20S β5 substrate

Cleavage products generated in the presence and absence of 500 nM of CFZ to knock out β5 activity were compared. The DMSO control reaction generated 197 cleavage sites after 1 h. In the presence of CFZ, the intensity of 69 of those cleavage products was reduced by 20-fold or more. Examples of a peptide that show this is shown in **Fig. 2A**, demonstrating how product formation accumulates over time in the non-inhibited assay and how it is not formed in the presence of CFZ. We generated a substrate specificity profile of the amino acids in the P4 to P4′ sites of these 69 cleavage sites and found a preference for hydrophobic amino acids at P4 and P1, K and T at P2, and W, N, V and Y at P3 (**Fig. 2B**). On the prime-side, R, L, F and P were most frequently found at P1′, P2′, P3′ and P4′, respectively. In general, there were more amino acids that were significantly enriched on the non-prime side so we synthesized the substrate, Ac-FNKL-amc which corresponds to amino acids enriched in the P4 to P1 positions. The turnover rate of this substrate was compared to that of the standard human β5 subunit, Suc-LLVY-amc at increasing concentrations. We show that cleavage activity with Ac-FNKL-amc is maximal at 25 µM and occurs at a ∼four-fold higher rate than with Suc-LLVY-amc at the same concentration (**Fig. 2C**).

**Figure 2.**
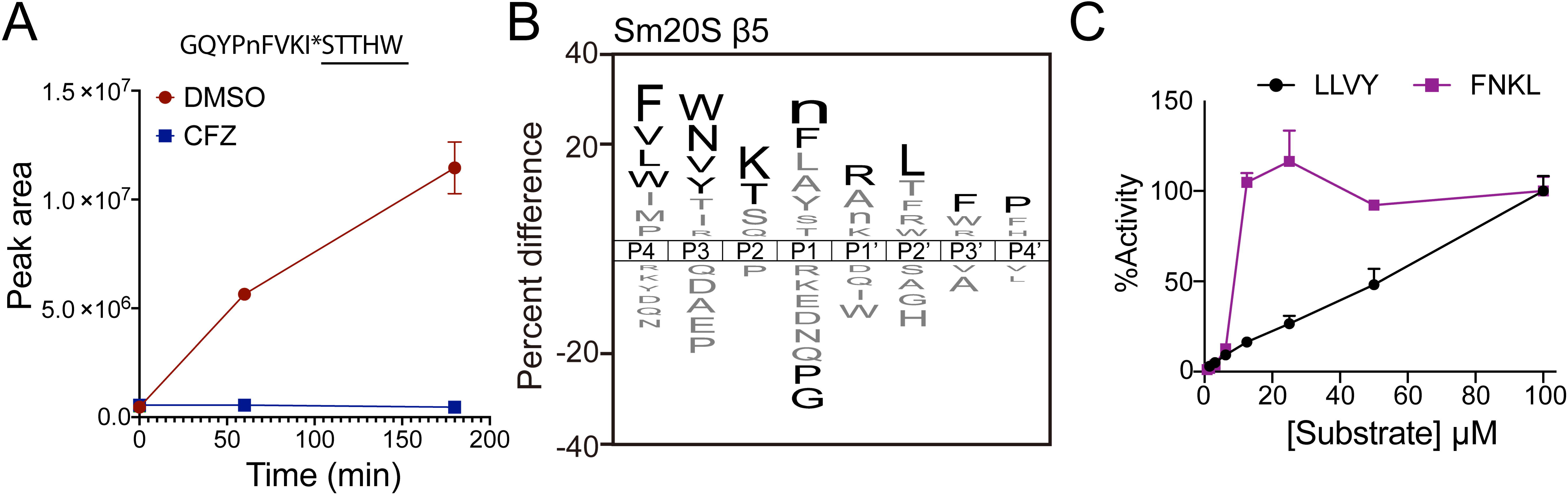
MSP-MS profiling of the Sm20S β5 catalytic subunit. **A**. An example of a single peptide cleaved by Sm20S in the MSP-MS assay. The abundance of cleaved peptides decreases in the presence of CFZ. **B**. IceLogo plot illustrating the substrate profile of all 69 peptide substrates that are cleaved by the putative β5 subunit**. C**. Comparison of the hydrolysis of Ac-FNKL-amc and Suc-LLVY-amc as a function of concentration. Fluorogenic assays were performed in technical triplicates while mass spectrometry-based assays were performed in technical quadruplicates.

### Development of a Sm20S β1 substrate

We next evaluated the MSP-MS cleavage products that were inhibited by 500 nM BTZ but uninhibited or weakly inhibited by 500 nM CFZ. We found 17 cleavage products that were inhibited by BTZ and not inhibited by CFZ. These cleavage products are likely, therefore, to be hydrolyzed by β1. An example of a substrate that is cleaved under these conditions is FDWWGNRSPLE*KnV (**Fig. 3A**). We attempted to synthesize several of these substrates that consisted of Ac-P4-P3-P2-P1-amc but had low yield and purity due to inefficient coupling of the Glu and Asp to the amc group. Therefore, we searched for protease substrates in commercial catalogs that contained E or D at P1 and that might be cleaved by Sm20S β1. We screened nine internally quenched caspase substrates and found that the sequence mca-VDQMDGW-K(dnp) was hydrolyzed by Sm20S. Importantly, this activity was inhibited by BTZ but not CFZ. This inhibition pattern was similar to that identified when using the substrate for the c20S β1 subunit, LLE-amc (**Fig. 3B**). When both substrates were evaluated over a concentration range of 2 to 500 μM, VDQMD*GW was cleaved with ∼2-fold higher efficiency between 16 and 250 μM (**Fig. 3C**).

**Figure 3.**
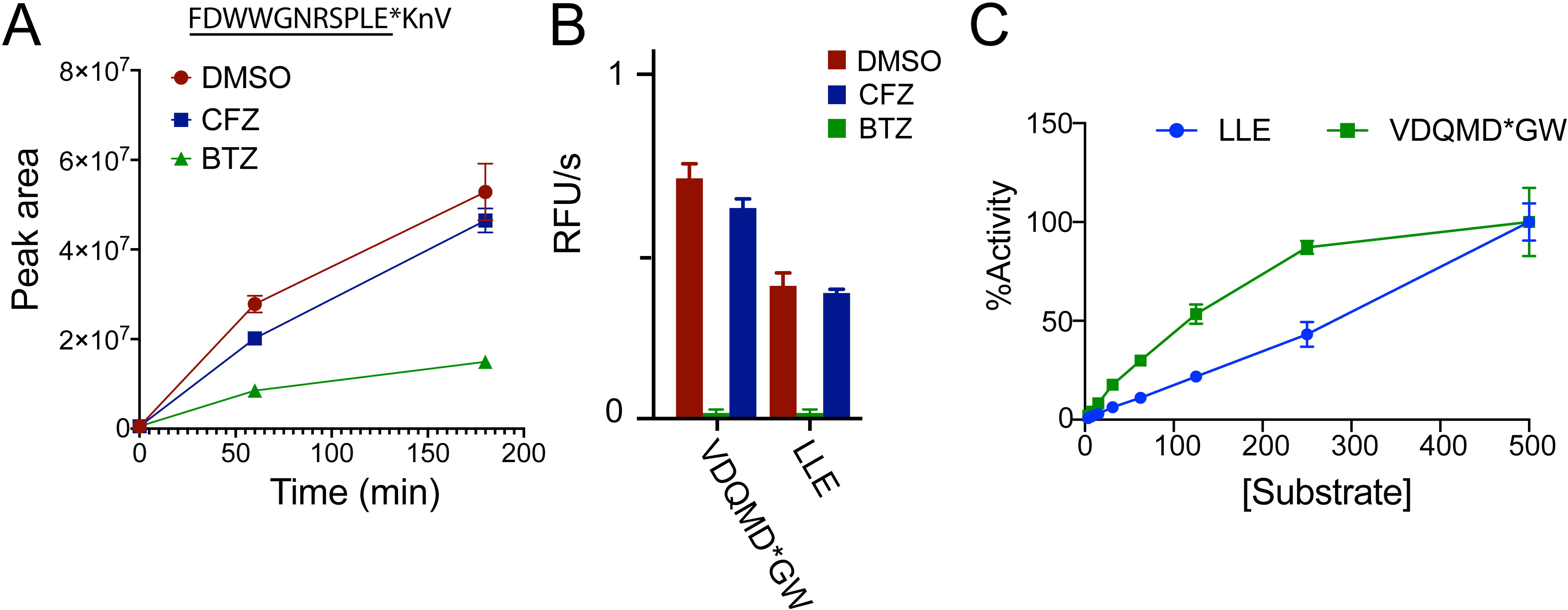
MSP-MS profiling of the Sm20S β1 catalytic subunit. **A**. An example of a single peptide cleaved by Sm20S in the MSP-MS assay showing the abundance of peptide as a function of the inhibitor used. **B**. Inhibition profile of Sm20S with a commercial caspase 3 substrate (mca-VDQMD*GW-K(dnp)) and Z-LLE-amc. **C**. Comparison of the hydrolysis of mca-VDQMD*GW-K(dnp) and LLE-amc as a function of concentration. Fluorogenic assays were performed in technical triplicates while mass spectrometry-based assays were performed in technical quadruplicates.

### Development of a Sm20S β2 substrate

We next evaluated the MSP-MS peptide cleavage profiles that were neither inhibited by 500 nM of BTZ or CFZ. We anticipated that many of these cleaved peptides are due to activity of the β2 subunit. We found 142 cleaved peptides that fit these criteria, exemplified by HWAFR*SRYHGPLAH (**Fig. 4A**). The overall profile indicated a high frequency of cleavage when R is present at P3, P1 and P1′, and K is present at P2 (**Fig. 4B**). We, therefore, synthesized an internally quenched substrate with the sequence mca-FnKRR-K(dnp) with which cleavage was predicted to occur between the two R residues. We included norleucine (n) at P3 instead of the preferred Arg residue to avoid cleavage at this alternate site. In our experience, extended pre-incubation of Sm20S with 10 μM of the irreversible inhibitor CFZ, inactivated Sm20S β2 (in addition to β5), while 10 μM of BTZ the reversible inhibitor only partially inhibited activity. Therefore, we used these two concentrations of BTZ and CFZ to evaluate if mca-FnKRR-K(dnp) is a β2 subunit. Sm20S was pre-incubated with 10 μM of CFZ and BTZ and then assayed with mca-FnKRR-K(dnp) and Z-LRR-amc. We showed that activity was inhibited by CFZ, similar to that noted with the canonical c20S β2 substrate, Z-LRR-amc (**Fig. 4C**). In addition, mass spectrometry confirmed the cleavage of mca-FnKRR-K(dnp) between the two Arg residues (**Fig. S3**). Cleavage of mca-FnKRR-K(dnp) and Z-LRR-amc and was then evaluated over a concentration range of 6.25 to 500 μM. For reasons as yet unclear, cleavage of mca-FnKRR-K(dnp) was biphasic, being most rapid at concentrations up to 50 μM, after which no further increase in activity was measured up to 500 μM (**Fig. 4C**). By contrast, cleavage of Z-LRR-amc steadily increased across the entire concentration range tested.

**Figure 4.**
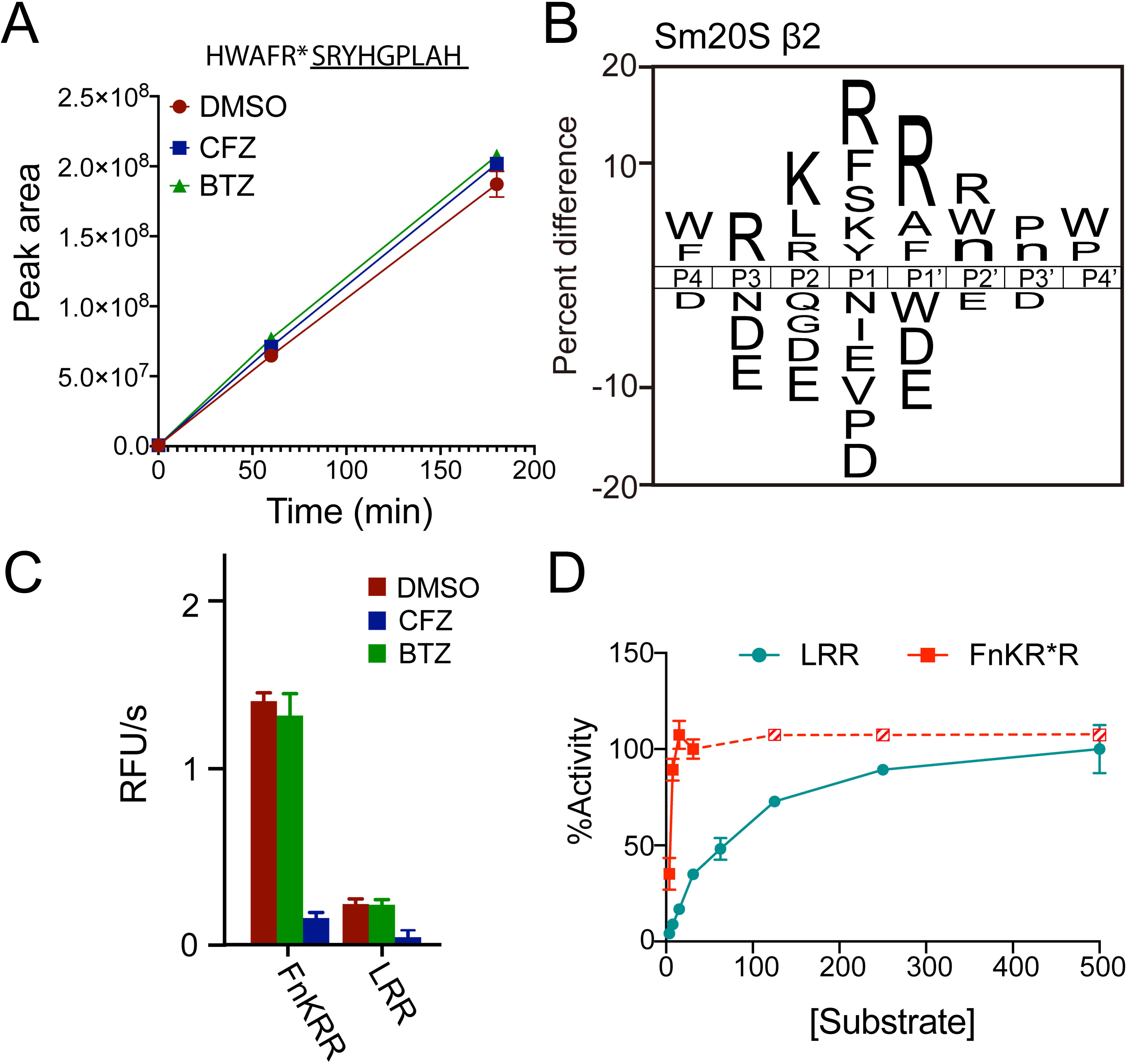
MSP-MS profiling of Sm20S β2 catalytic subunit. **A.** An example of three peptides cleaved by Sm20S in the presence of 500 nM BTZ and CFZ when compared to vehicle control (DMSO). **B.** iceLogo plot illustrating the substrate profile of 142 peptide substrates that are cleaved. **C.** Cleavage of mca-FnKRR-K(dnp) and Z-LRR-amc in the absence and presence of 10 μM of BTZ and CFZ. **D.** Comparison of the hydrolysis of mca-FnKRR-K(dnp) and Z-LRR-amc as a function of concentration. Fluorogenic assays were performed in technical triplicates while mass spectrometry-based assays were performed in technical quadruplicates.

### Testing new substrates with single-step enriched Sm20S from cell lysates

The isolation of Sm20S to homogeneity is costly and labor-intensive due to the difficulty in scaling up the production of *S. mansoni* which must be harvested from vertebrate animals. Therefore, armed with our three new substrates, we evaluated Sm20S activity in *S. mansoni* protein extract that had only been subjected to ammonium sulfate precipitation, the first step of the three-step purification process. Pepstatin, E-64, and AEBSF were included in the assay buffer to prevent substrate cleavage by *S. mansoni* aspartyl, cysteine and serine proteases, respectively in the single-step enriched Sm20S. ^59–61^ We show that β1, β2 and β5 activities were all detected in the enriched extract and that the inhibition profile using BTZ and CFZ matched that of the purified enzyme (**Fig. 5A**). These studies confirm that the single-step enriched Sm20S is sufficient to specifically evaluate the β1, β2 and β5 activities.

**Figure 5.**
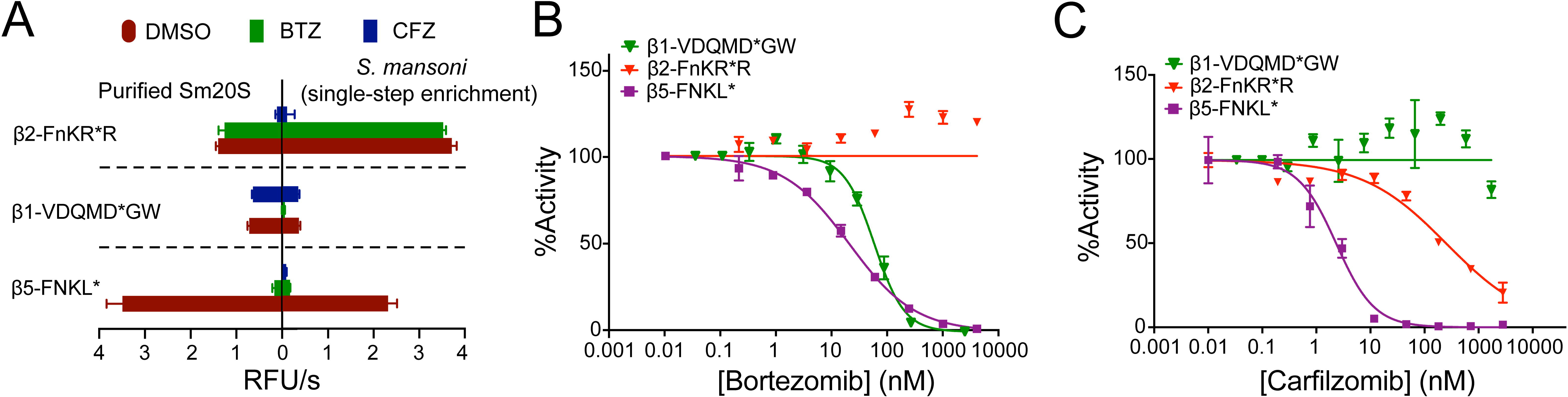
Evaluation of Sm20S substrates with pure Sm20S and single-step enriched Sm20S. **A**. Comparison of the activities of purified Sm20S and the single-step enriched Sm20S. **B**. Comparison of the hydrolysis of the three β subunit specific substrates by single-step enriched extract in the presence of different concentrations of BTZ. **C**. Comparison of the hydrolysis of the three β subunit-specific substrates in the presence of different concentrations of CFZ. Data are shown as means ± SD of technical triplicates.

We next determined whether the potency of BTZ and CFZ could be directly determined in the single-step enriched Sm20S. BTZ has an IC_50_ value of 49.84 ± 0.24 nM for the β1 subunit (**Fig. 5B**), whereas CFZ was not inhibitory (**Fig. 5C**). Using the β2 substrate, BTZ did not inhibit this activity, whereas CFZ had an IC_50_ of 288.0 ± 90 nM. Using the β5 substrate, BTZ and CFZ had IC_50_ values of 17.97 ± 2.05 nM and IC_50_ = 2.495 ± 0.51 nM, respectively. These data demonstrate that proteasome inhibition assays can be performed using the single-step enriched Sm20S.

### The subunit specificity of 20S proteasomes from three *Schistosoma* species

The new β1, β2 and β5 substrates were designed based on the specificity profile of the 20S from *S. mansoni*, which is a model laboratory schistosome. New anti-schistosomal drugs should be active against *S. mansoni* and the other two medically important schistosomes, *S. haematobium* and *S. japonicum* ^51, 52^. In this regard, it is important to understand whether the substrates developed for Sm20S can be also used for the other two species. We first normalized proteasome activity in extracts of all three schistosome species using the proteasome inhibitor probe, Me4BodipyFL-Ahx3Leu3VS ^62^. This probe labeled three subunits of c20S with the bottom two bands co-migrating to yield one strongly fluorescent product (**Fig. 6A**). The probe apparently labelled two of the three β subunits in each of the schistosome extracts.

**Figure 6.**
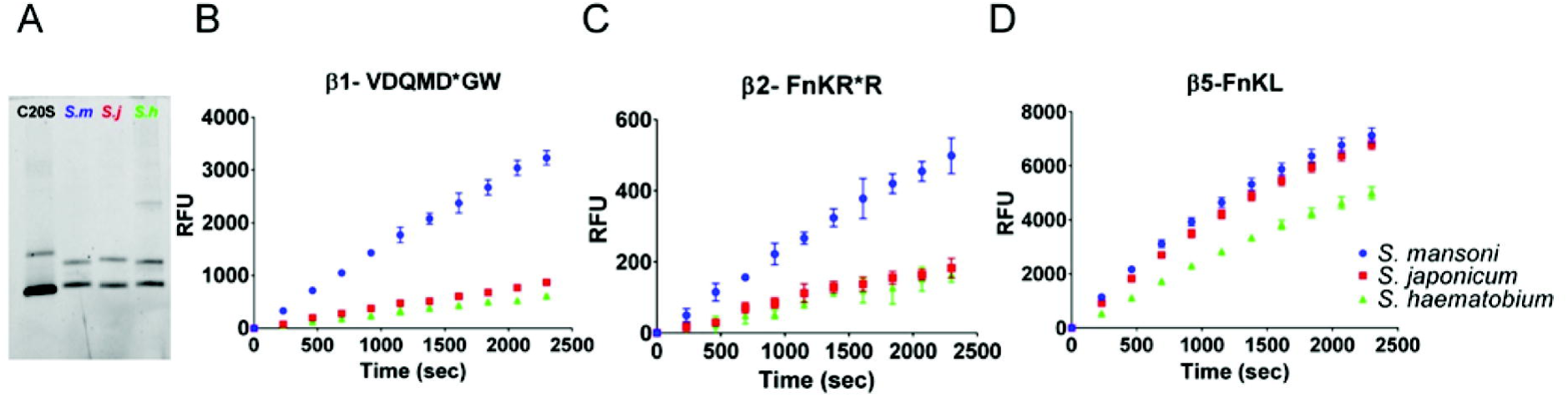
Comparison of the 20S activities from three *Schistosoma* species with the newly-designed substrates. **A**. *Schistosoma* spp. proteasome detection using the fluorescent Me4BodipyFL-Ahx3Leu3VS probe. This probe was used to normalize proteasome amount in advance of the in-solution fluorogenic substrate assays. 2.5 μg *S. haematobium* lysate and 5 μg *S. japonicum* and *S. mansoni* lysates were used in each lane and c20S was used as a control. **B-D**. *Schistosoma* spp. β1, β2 and β5 subunit activities in the presence of the MCA-VDQMD*GW-K-(DNP)-NH_2_, MCA-FnKRR-K-(DNP)-NH_2_ and Ac-FNKL-amc, respectively. Data are shown as means ± SD of technical triplicates.

Having used the probe to normalize for proteasome activity, we evaluated the three subunit-specific substrates across the single-step enriched 20S proteasomes from the three schistosome species. The specific activity using the β1 and β2 substrates was two-to three-times higher for S*. mansoni* than for *S. haematobium* and *S. japonicum* (**Fig. 6, B-D**), whereas the activities measured using the β5 substrate were similar for the three species. Schistosome lysates were pre-incubated with 500 nM BTZ or CFZ to test their cross-species inhibitory effects. Similar inhibition profiles were recorded (**Fig. 7**). BTZ preferentially inhibited β1 and β5, whereas CFZ inhibited β5 and β2 more than β1. In addition, we tested MZB, a marine natural product that is in Phase 3 clinical trials for the treatment of glioblastoma. ^26^ MZB decreased the activity measured with each of the three substrates, with complete inactivation of β5, 66 to 48% reduction of β2, and 61 to 34% reduction of β1 (**Fig. 7**). Overall, therefore, the inhibition responses by each of the three subunits for each of the three species is similar. It seems that we can, with reasonable confidence, extrapolate data acquired for future proteasome inhibitors from one schistosome species to the other two.

**Figure 7.**
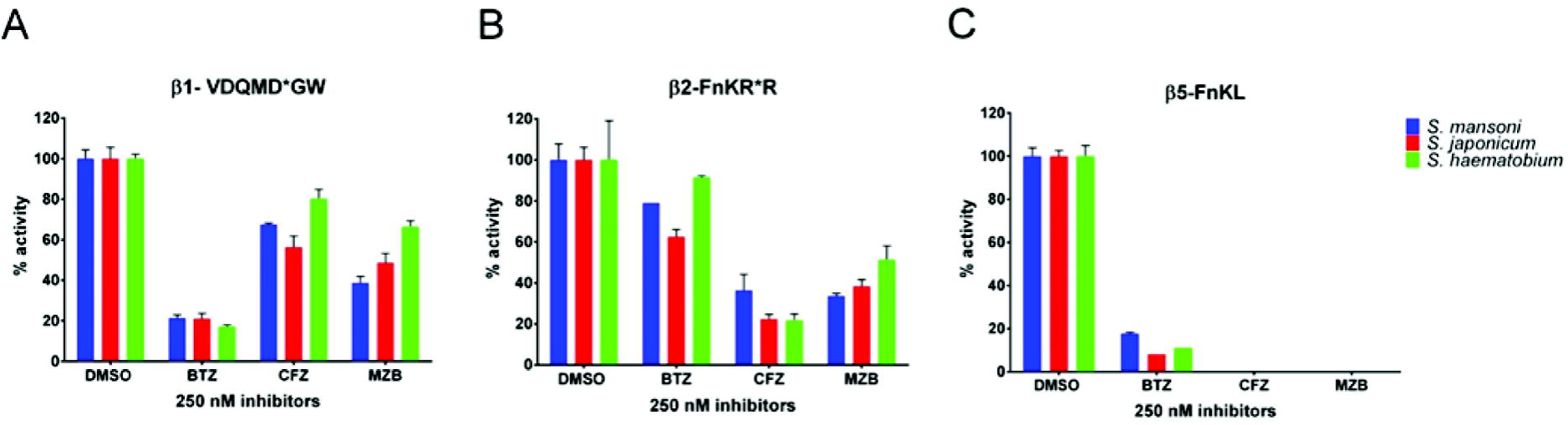
Comparison of proteasome inhibition across the three *Schistosoma* species. Proteasome inhibitors were pre-incubated at 500 nM with single-step enriched 20S enzymes from the three *Schistosoma* species. **A-C**. Inhibition profiles using the β1-, β2- and β5-subunit -specific substrate. Data are shown as means ± SD of technical triplicates.

We have previously shown that carmaphycin-17, an analogue of the marine natural product, carmaphycin B, demonstrated potent anti-schistosomal activity. ^21^ Using the subunit-specific substrates, we evaluated the potency of carmaphycin-17 at 1,000, 250 and 50 nM. We show that at 1 μM, carmaphycin-17 decreased Sm20S and Sh20S β1 activities by 20%, whereas the Sj20S β1 activity was somewhat more sensitive with a 47% reduction (**Fig. 8A**). For the β2 subunits, carmaphycin-17 was similarly potent with a ∼60% inhibition at 1 μM (**Fig. 8B**). Last, carmaphycin-17 was most potent against the β5 subunits with complete inhibition at 1 μM (**Fig. 8C**). Taken together, these data show that each of the three subunits in each of the three medically important species responds similarly to carmaphycin-17, with the exception of Sj20S β1. When combined with the data from **Fig. 7**, it seems that we can, with reasonable confidence, extrapolate data acquired for future proteasome inhibitors from one species to the other two.

## Discussion

For over 40 years, treatment and control of schistosomiasis have relied on one drug, praziquantel, and concerns regarding the emergence and establishment of resistance, in addition to the drug’s pharmaceutical and pharmacological limitations, have spurred the search for alternate chemotherapeutics. Among several molecular drug targets to have been considered over the years, the recent clinical success of targeting the *Plasmodium* and *Leishmania* 20S proteasome with small molecule inhibitors (cited above) has focused our attention on the schistosome 20S proteasome ortholog.

The proteasome is a large multi-subunit, ATP-dependent proteolytic complex that regulates several cellular processes, including normal protein turnover and degradation of misfolded proteins. Inhibitors of the human constitutive c20S, *e.g.,* CFZ and BTZ, are key drugs for treatment of blood cancers, and, more recently, Zetomipzomib (KZR-616) is one of several new inhibitors of the i20S that are in clinical trials for treatment of autoimmune diseases and immune-mediated disorders. ^63^ Anti-cancer proteasome inhibitors that target Sm20S were shown to decrease worm motility *in vitro* and kill the parasite.^21^ Also, inhibition of 19S proteasome deubiquitinating activity in *S. mansoni* induced a modest reduction in egg production *in vitro*, decreased viability, and were eventually cidal ^22^. Taken together, these data suggest that the schistosome proteasome is a drug target of interest for the potential treatment of schistosomiasis, and support the further characterizing of the cleavage specificities of the individual catalytic β subunits of Sm20S, information that would underpin a campaign to develop Sm20S-selective inhibitors.

Each of the 20S catalytic β subunits has a distinct substrate specificity that varies between species ^64^. Fluorogenic substrates such as Z-LLE-amc, Z-LRR-amc, and Suc-LLVY-amc that were developed for the human c20S β1, β2, and β5 subunits, respectively, are also cleaved by the respective subunits of parasite proteasomes ^32, 65^. However, as we have discovered here using MSP-MS-directed substrate design, these c20S substrates are not optimal for the Sm20S β1, β2, and β5 subunits, and we were able to optimize a new substrate for each subunit that performed two-to eight-fold better. These rationally optimized β subunit-selective substrates for Sm20S, therefore, should prove valuable in the design of inhibitors targeting each β subunit or combinations of β subunits. As a case in point, our MSP-MS studies suggest that the preferentially cleavage of the sequence RKR*R (where * is the cleavage point) by Sm20S β2 (Fig. 3B) is different from the cleavage preference of c20S β2 subunit which indicates that positively charged residues at P3, P2 and P1′ are not so enriched ^58^. Accordingly, our future inhibitor design efforts will focus on exploiting this and other differences between c20S and Sm20S.

We also developed a single-step enrichment of Sm20S activity from crude extracts using ammonium sulfate, which, when combined with the new specific substrates, allowed for a quicker evaluation of inhibitor specificity and action. Isolation of Sm20S, although initially required to define the cleavage specificities of the individual catalytic β subunits and facilitate substrate design, is no longer needed with the new specific substrates in hand. This will save time and the materiel required (including vertebrate animals) to isolate limiting amounts of pure enzyme. Our findings of similar inhibition response profiles for the three catalytic β subunits across the three medically important schistosome species with either the anticancer inhibitors or our in-house carmaphycin B analog, initially suggest that an inhibitor design and development campaign focused on Sm20S could quickly translate to the two other orthologous targets as well. This woud be in line with the preferred target product profile for schistosomiasis ^51, 52^ which calls for drugs that are active against all schistosome species, or at least the two species predominating in Africa where >90% of the global burden of schistosomiasis exists. Also, the ability to focus on the most experimentally tractable of the three species, *i.e., S. mansoni*, to develop β subunit-selective inhibitors will save time and money, especially important in the context of diseases of poverty for which the financial support for drug development is constrained.

In summary, MSP-MS has helped characterize the substrate specificities of the three catalytic β subunits of Sm20S which subsequently directed the design of optimized substrate for each subunit. These substrates were validated in worm extracts enriched for Sm20S in a single step, eliminating the need for lengthy and costly purifications of the enzyme. Finally, the optimized substrates demonstrated that the 20S proteasome is similar across the three major *Schistosoma* species, suggesting that a program to develop Sm20S inhibitors will likely yield compounds that are broadly schistosomicidal.

## Funding

The research was supported by NIH awards R21AI133393 and R21AI171824 to AJO and CRC and R01AI158612 and R21AI146387 to AJO. The content is solely the responsibility of the authors and does not necessarily represent the official views of the National Institutes of Health. PF received funding from the Program for Research and Mobility Support of Starting Researchers from the Czech Academy of Sciences (MSM200551901) and the European Union’s Horizon 2020 research and innovation program under the Marie Skłodowska-Curie grant agreement No. 846688, ProTeCT.

---

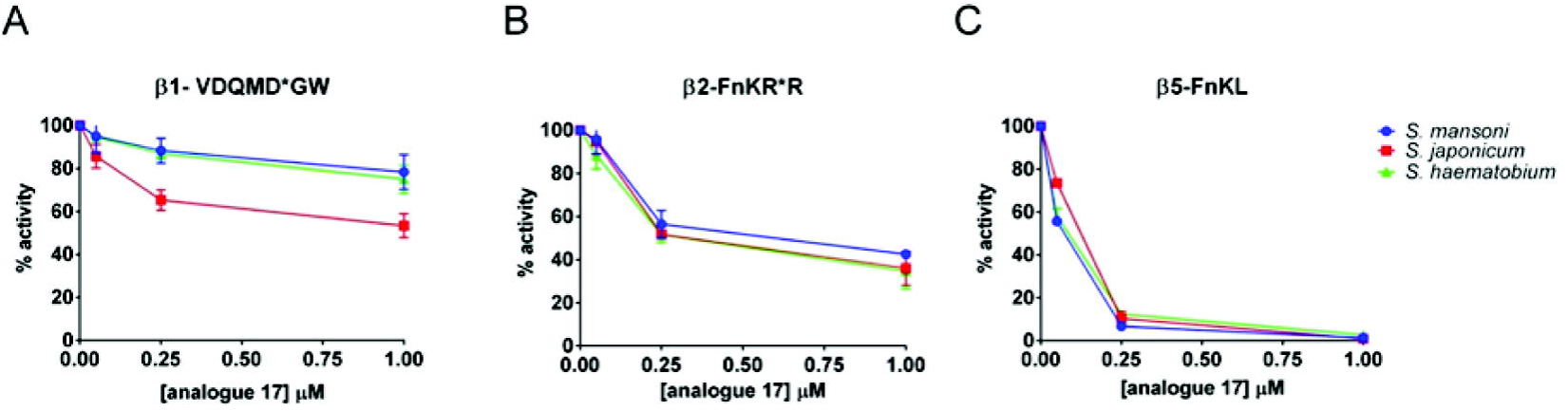

## Notes

### Competing Interest Statement

The authors have declared no competing interest.

